# NADPH oxidase inhibitor enhances brain resilience in Alzheimer’s disease by reducing tauopathy and neuroinflammation

**DOI:** 10.1101/2025.03.01.640956

**Authors:** Jihyeon Lee, Seunghwan Sim, Yinglan Jin, Hyejun Park, Eun Young Byeon, Su Jin Kim, Sujin Yun, Hye Eun Lee, Da Un Jeong, Jung Min Suh, In Hye Lee, Ho-Young Lee, Yongseok Choi, Yun Soo Bae

## Abstract

Alzheimer’s disease associates closely with activation of NADPH oxidase (Nox) isozymes. We identified that CRB-2131, a novel oxadiazole derivative, potently suppresses Nox isozymes. It inhibits reactive oxygen species production (ROS) by hippocampal neuronal and microglial cells and reduces microglial activation. Prophylactic (starting at 3.5 months of age) and therapeutic (starting at 6 months of age) oral administration with CRB-2131 for 10 weeks in 5XFAD mice reduced hippocampal superoxide levels, lipid peroxidation, Tau phosphorylation, and neuroinflammation. Prophylactic and therapeutic CRB-2131 treatment of 5XFAD mice restored their impaired cognition as shown by the novel-object recognition, Y-maze, and Morris water-maze tests. CRB-2131 treatment increased mature neurons, reduced apoptotic mature neurons, and elevated immature neurons in the hippocampus. Positron-emission tomography/computed-tomography imaging confirmed that CRB-2131 stimulated neuronal regeneration. CRB-2131 suppresses brain oxidation, tauopathy, and neuroinflammation, thereby preventing mature neuron death and promoting neuron regeneration. Ultimately, this fosters a resilient brain and protects cognition.

## Introduction

Alzheimer’s disease (AD) is a highly complex disease and thus presents a formidable challenge in the field of neurodegenerative disorders. It is characterized by multiple neuropathological hallmarks, including amyloid-β (Aβ) plaques, neurofibrillary tangles, and significant neuronal loss^1, 2, 3^. These pathological events in AD appear to be driven by oxidative stress and neuroinflammation^4, 5, 6^. Several lines of evidence suggest that oxidative stress may promote the abnormal accumulation and aggregation of Aβ protein, which then spreads throughout the central nerve system (CNS) parenchyma. The Aβ protein in turn activates the brain-resident macrophages and microglia, which produce pro-inflammatory mediators^7, 8^. Ultimately, this and other neuroinflammatory pathways lead to tauopathy. However, there is also evidence that neuroinflammation may conversely promote oxidative stress, and that these two pathological entities dovetail in the development and progression of AD.

Recent research has yielded intriguing results regarding the progression of cognitive decline in elderly individuals. In a longitudinal study involving 1,325 cognitively normal older adults, amyloid PET and tau PET scans were used to monitor these individuals over a 3.5-year period^9^. The participants were classified into four distinct groups: (i) A^-^T^-^, with both β-amyloid (A) and tau (T) negative; (ii) A^+^T^-^, with β-amyloid positive but tau negative; (iii) A^+^T_MTL_ ^+^, with β-amyloid positive and tau positivity localized in the medial temporal lobe; and (iv) A^+^T ^+^, with β-amyloid positive and tau positivity in the neocortex. The findings indicated that the A^+^T_MTL_^+^ and A^+^T ^+^groups exhibited a significantly higher risk of progressing to mild cognitive impairment (MCI) or dementia, along with a more rapid decline in cognitive function. These results underscore the correlation between tau pathology and cognitive decline in clinical settings, suggesting that tau pathology may be more closely associated with cognitive impairment, a hallmark of Alzheimer’s disease^9^.

Oxidative stress plays a critical role not only in the early stages of tauopathy by promoting abnormal tau phosphorylation and accumulation, which exacerbates neuronal damage, but also in inducing mitochondrial dysfunction and NADPH oxidases (Nox) activation due to such aberrant tau accumulation. This, in turn, generates additional reactive oxygen species (ROS), thereby intensifying oxidative stress. This process can create a vicious cycle that accelerates the progression of tauopathy^10^. Consequently, targeting Nox, which are responsible for ROS production, presents a highly competitive and promising therapeutic strategy for mitigating the pathological cascade wherein Aβ exacerbates toxic tau, potentially offering new avenues for treating dementia. Recent research suggests that the Nox isozymes in the brain play pivotal roles in AD by shaping both oxidative stress and neuroinflammation. The Nox enzymes regulate the production of reactive oxygen species (ROS)^11, 12^. To date, seven Nox isozymes [Nox1, Nox2, Nox3, Nox4, Nox5, Dual oxidase 1 (Duox1) and Dual oxidase 2 (Duox2)] have been identified in various human cell types^13^. Neurons express Nox1, Nox2 and Nox4, while Nox2 is exclusively expressed in microglia and astrocytes^14^. Several recent studies suggest that Nox2 on neurons and microglia and Nox4 on neurons may promote AD-like disease by inducing oxidative stress and neuroinflammation. First, Tu et al. found that inhibiting or deleting Nox2 prevented inflammation-elicited neuronal production of ROS in vitro ^14^. Second, Luengo et al. showed that neuronal Nox4 deficiency in a humanized mouse model of tauopathy reduced Tau-induced neurotoxicity and suppressed cognitive decline^13^.

These observations together suggest that therapeutically targeting the interplay between various Nox isozymes and oxidative stress and neuroinflammation could mitigate AD progression^15^. This approach is of particular interest because it targets the underlying pathogenic mechanisms of AD rather than merely addressing its symptoms. We show here that a promising candidate Nox inhibitor is the novel oxadiazole derivative CRB-2131: our experiments with 5XFAD mice, a well-established animal model of AD, showed that CRB-2131 not only markedly curtailed ROS production in the brain, it also ameliorated tauopathy and neuroinflammation, thereby promoting the regeneration and repair of neural tissue and the cognitive functions of the mice.

## Results

### The Nox inhibitor CRB-2131 inhibits Aβ-induced neuronal and microglial ROS production

Given the studies showing that brain Nox isozymes associate with AD^16, 17, 18, 19^, we subjected a chemical library containing 45,000 compounds to a high-throughput Nox inhibitor-screening assay to identify chemical compounds that could inhibit Nox activity. Thus, purified Drosophila membranes expressing human Nox1, Nox2, Nox4, Nox5, Duox1, or Duox2 were incubated with each compound and their ROS production was monitored by determining the oxidation of lucigenin, which was evident as chemiluminescence^20^. The screening revealed oxadiazole analogues that inhibited Nox isozymes. We then synthesized 215 compounds by modifying the parent molecule and examined the relationship between the structure of the compound and its ability to inhibit Nox. This led to the identification of the oxadiazole derivative 4-fluoro-N-(5-phenyl-1,3,4-oxadiazol-2-yl)-3-(trifluoromethyl) benzamide, which was designated CRB-2131 (Fig. 1A). The potential of this compound as a drug candidate was suggested by its *in vitro* Nox-inhibitory activity, as measured with the Nox-bearing *Drosophila* membrane system described above: the IC_50_ values of all Nox isozymes ranged from 0.2 to 0.3 μM (Fig. 1B).

**Figure 1.**
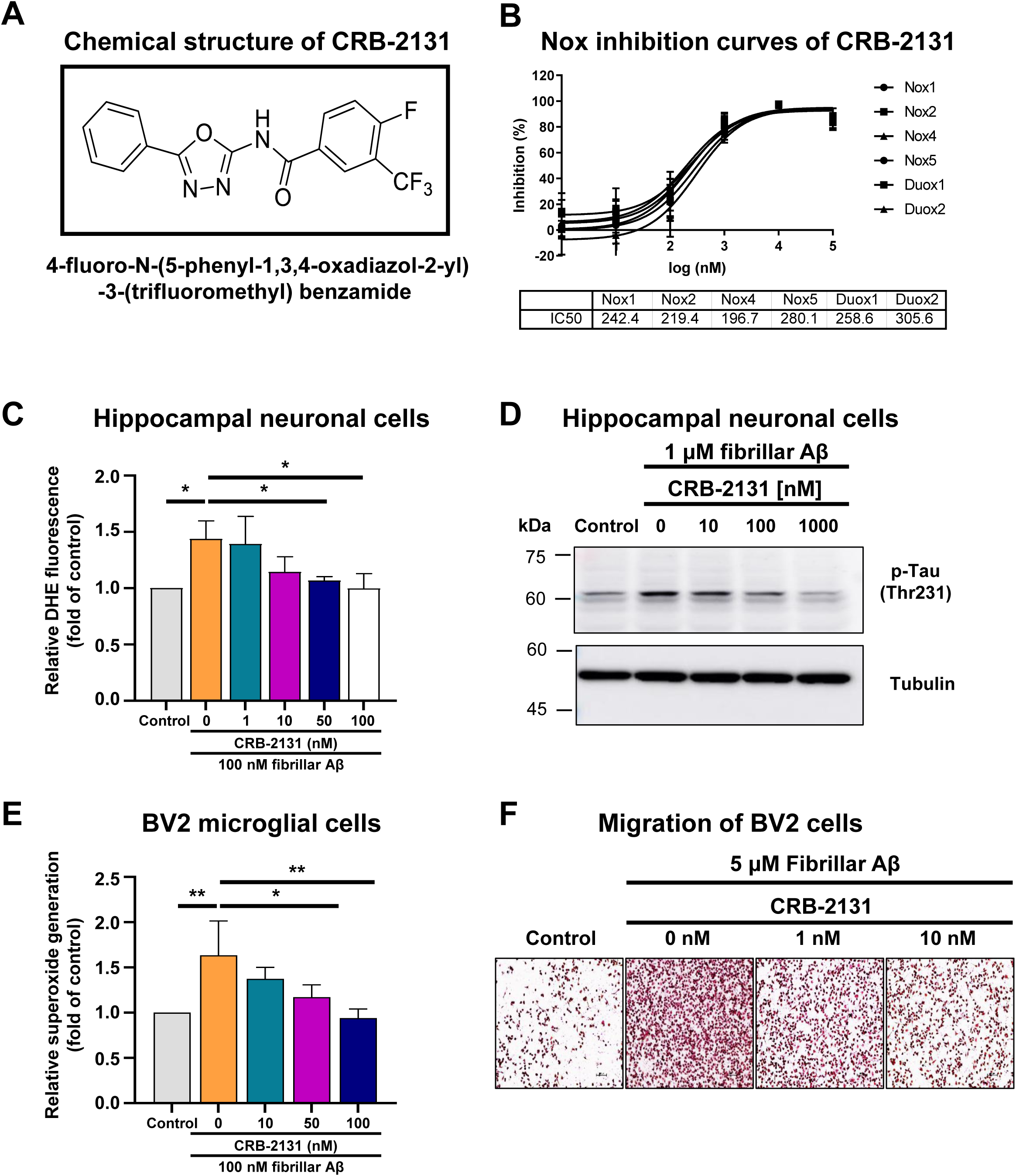
Chemical structure of CRB-2131 and its effect on neuronal and microglial ROS production, Tau phosphorylation, and neuroinflammation. (A) Chemical structure of CRB-2131. B) Concentration-dependent Nox inhibition curves of CRB-2131. Drosophila membranes specifically overexpressing hNox1, hNox2, hNox4, hNox5, hDuox1, and hDuox2 were subjected into ROS measurement with Diogene. The IC_50_ value of CRB-2131on hNox1, hNox2, hNox4, hNox5, hDuox1, and hDuox2. C and D) Dose-dependent inhibitory activity of CRB-2131 on Aβ-mediated ROS generation (C) and Tau phosphorylation (D) in HT-22 cells. (N=3, mean ± SD, *p<0.05). E) Dose-dependent inhibitory activity of CRB-2131 on Aβ-mediated ROS generation in BV2 microglial cells. (N=3, mean ± SD, *p<0.05, **p<0.01). F) Transwell migration assay. BV2 microglial cells were seeded into upper chamber and stimulating medium containing Aβ was added to the lower chamber. After 16hrs, the migrated cells were stained with hematoxylin/eosin and counted. Representative images for each group.

We next asked whether CRB-2131 could inhibit Aβ-induced activities of neurons and microglia that associate closely with AD pathogenesis and progression^16, 18^. Indeed, CRB-2131 potently inhibited the Aβ-induced (i) production of ROS by HT-22 mouse hippocampal neuronal cells (IC_50_=5.98 nM) (Fig. 1C), (ii) phosphorylation of Tau in the HT-22 neuronal cells (Fig. 1D), (iii) ROS production by BV2 microglial cells (IC_50_=17.1 nM) (Fig. 1E), (iv) expression of TNF-α and IL-1β by BV2 microglial cells (fig. S1, A and B), and (v) BV2 microglial-cell migration (IC_50_=0.96 nM) (Fig. 1F and Fig. S1C).

Next, we investigated whether CRB-2131 can directly scavenge ROS in vitro. Thus, 1.0 and 10 μM of CRB-2131 was incubated with H_2_O_2_ and lucigenin: the oxidation of lucigenin by H_2_O_2_ generated luminescence, which served as a measure of H_2_O_2_ concentration. We added hydrogen peroxide in the presence or absence of CRB-2131 (1.0 and 10 μM) and then measured changes in H_2_O_2_ concentration. The positive control (the ROS scavenger N-acetyl cysteine) effectively blocked H_2_O_2_-induced lucigenin oxidization but CRB-2131 had no ability to scavenge ROS (Fig. S2).

### CRB-2131 treatment suppresses brain ROS and pTau levels in 5XFAD mice

To determine the ability of CRB-2131 to block AD pathology in animal model, male 3.5-month-old 5XFAD mice were given a single 10 mg/kg (mpk) dose of CRB-2131 orally. The brains of 5XFAD mice overexpress human amyloid-β precursor protein 695 (APP) and human presenilin-1 that bear multiple familiar AD mutations. Consequently, these mice start developing amyloid plaques and neurodegeneration at 3.5 months of age. Based on the pharmacokinetic C_max_, brain-to-plasma ratio (Fig. S3), and the molecular weight of CRB-2131, the maximal concentration of CRB-2131 in the brain of 5XFAD mice after a single 10 mpk CRB-2131 dose was 270 nM, which exceeds the IC_50_ of CRB-2131 in terms of HT-22-cell ROS production (IC_50_=5.98 nM), BV2-cell ROS production (IC_50_=17.1 nM), and BV2-cell migration (IC_50_=0.96 nM) (Fig. 1, E and F). The peak concentration of CRB-2131 in the brain was reached at 30 minutes after administration (Fig. S3).

High ROS levels in the brain associate strongly with AD and 5XFAD mice have high brain ROSs^21, 22^. To test whether CRB-2131 can reduce the formation of these ROS, we used a prophylactic treatment regimen: thus, 3.5-month-old 5XFAD mice were treated *q.d.* (*quaque die*, once a day) and *P.O.* (*per os*, oral administration) with 1, 3, or 10 mpk CRB-2131 for 10 weeks and the now-6-month-old mice were sacrificed (Fig. S4A). The harvested brain tissue was then stained with dihydroethidium (DHE) dye, which detects superoxide. Compared to untreated WT mice, the vehicle-treated 5XFAD mice demonstrated elevated ROS levels in the dentate gyrus (DG) of the brain, but CRB-2131 treatment decreased these ROS levels in a dose-dependent fashion in DG (Fig. 2, A and B).

**Figure 2.**
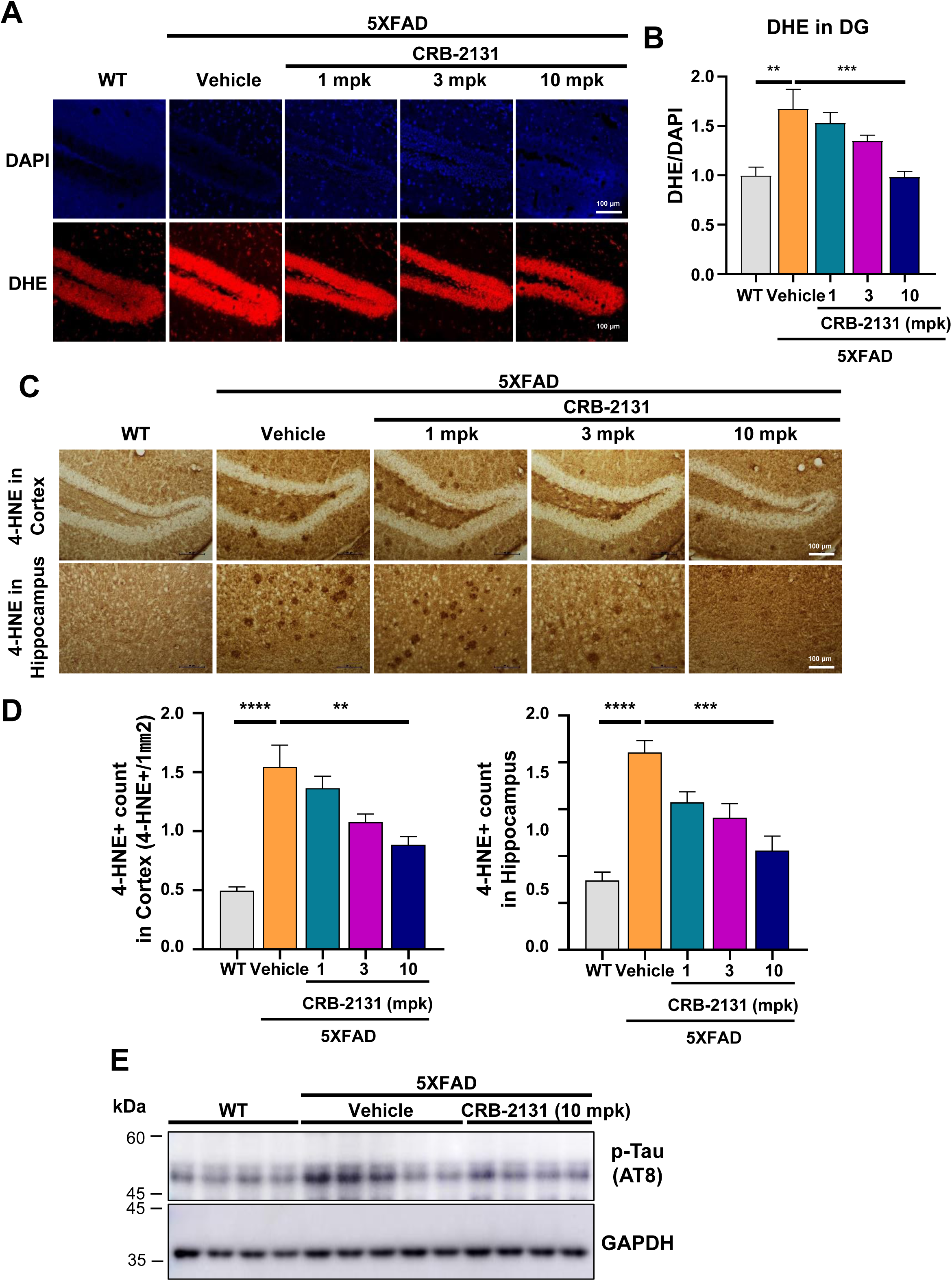
CRB-2131 inhibits ROS generation and Tau phosphorylation in 5XFAD mice. (**A** and **B**) Ability of prophylactic CRB-2131 (1, 3, or 10 mg/kg of CRB-2131, *q.d.*, *P.O.*, and 10 weeks) to suppress ROS generation in the DG of 5XFAD mice, as determined by DHE staining after sacrifice. The mpk means mg/kg. (A) Representative images. (B) Quantification of DHE levels. (**C** and **D**) Ability of prophylactic CRB-2131 to inhibit 4-HNE levels in the brain of 5XFAD mice. (C) Representative images. (D) Quantification of 4-NHE levels (**E**) Ability of prophylactic CRB-2131 to inhibit Tau phosphorylation (pTau) in the brain of 5XFAD mice. All quantitative data in this figure are shown as mean ± SD. **p<0.01, ***p<0.005, as determined by one-way ANOVA followed by Turkey’s post-hoc test.

Brain tissue is rich in polyunsaturated fatty acids. This lipid-rich environment is highly susceptible to oxidative stress, which can lead to lipid peroxidation. This is also observed in the brain of 5XFAD mice^23, 24^. Thus, we examined the effect of CRB-2131 on the 4-hydroxynonenal (4-HNE) adducts, which are a marker of brain lipid oxidation in AD^25^.

Immunohistochemistry with an antibody against 4-HNE showed that the vehicle-treated 5XFAD mice had higher 4-HNE levels in the brain than untreated WT mice, and that this was attenuated by prophylactic CRB-2131 treatment in a dose-dependent manner (Fig. 2, C and D). Tau phosphorylation associates strongly with AD^26, 27^. We found that the vehicle-treated 5XFAD mice had more Tau phosphorylation in their brain than untreated WT mice, and that 10 mpk CRB-2131 suppressed this (Fig. 2E).

### CRB-2131 suppresses Tau phosphorylation and neuronal inflammation in 5XFAD mice

Next, we assessed the in vivo mechanism by which the Nox inhibitor CRB-2131 improves 5XFAD mouse cognition. We first found that prophylactic CRB-2131 treatment (3.5-month-old mice, 1, 3, or 10 mpk, *q.d.*, *P.O.*, and 10 weeks) did not regulate Aβ aggregation in the brains of 5XFAD mice (Fig. 3A and Fig. S5A). However, it did reduce the pTau levels in the neurofibrillary tangles in the cortex and hippocampus of these mice in a dose-dependent manner (Fig. 3A and Fig. S5B and S5C). This is significant because Tau phosphorylation is observed in both 5XFAD mice and AD patients and pTau levels correlate with cognitive decline in AD patients^26, 27^. These results indicate that CRB-2131 inhibits pTau level in brain of 5XFAD mice (Fig. 2E and Fig. 3A).

**Figure 3.**
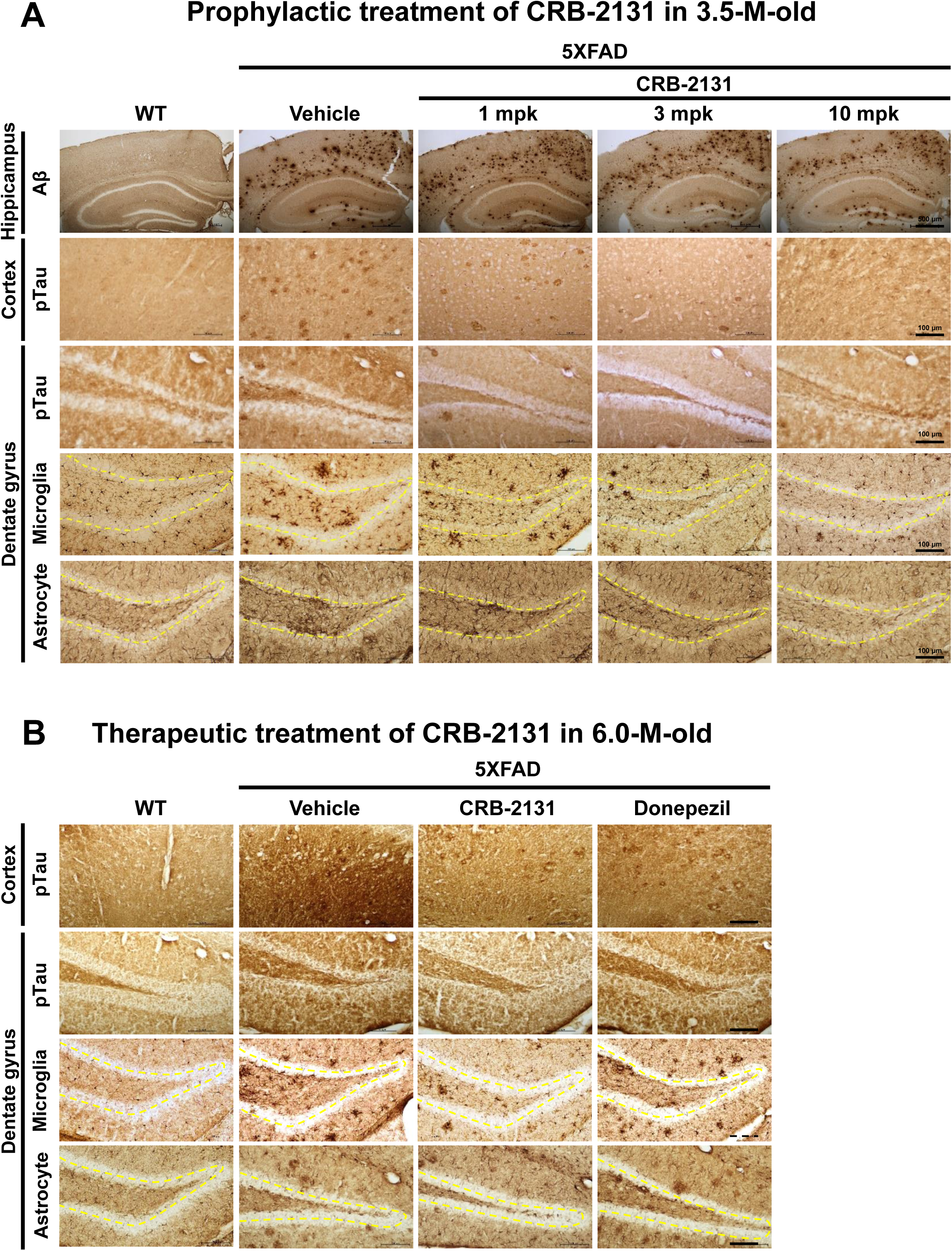
Prophylactic and therapeutic CRB-2131 suppresses Tau phosphorylation and neuronal inflammation in 5XFAD mice. **(A)** Prophylactic CRB-2131-treated 5XFAD mice (1, 3, or 10 mg/kg of CRB-2131, *q.d.*, *P.O.*, and 10 weeks) were subjected to immunohistochemistry of brain Aβ, phosphorylated Tau (pTau) in the cortex and the hippocampus, Iba1^+^ for microglial cells, and GFAP^+^ for astrocytes. Immunohistochemistry of Aβ, pTau in cortex and DG, Iba1 in DG, and GFAP in DG of 5XFAD mice with or without CRB-2131 dose-dependent administration. The mpk means mg/kg. **(B)** Therapeutic CRB-2131-treated 5XFAD mice (6.0-month-old mice, 10 mg/kg of CRB-2131, *q.d.*, *P.O.*, and 10 weeks) were subjected to immunohistochemistry of brain phosphorylated Tau (pTau) in the cortex and the hippocampus, Iba1^+^ for microglial cells, and GFAP^+^ for astrocytes.

We then found with immunohistochemistry of the brain that prophylactic CRB-2131 treatment (3.5-month-old mice, 1, 3, or 10 mpk, *q.d.*, *P.O.*, and 10 weeks) improved the neuroinflammation in 5xFAD mice: they demonstrated significantly fewer GFAP^+^ astrocytes and Iba-1^+^ microglia in the DG of the hippocampus than the vehicle-treated 5XFAD mice (Fig. 3A and Fig. S5D and S5E). A CRB-2131 dose-dependent effect was also observed (Fig. 3A). Since AD associates with increased activation of GFAP^+^ astrocytes and Iba-1^+^ microglia and their migration into the DG^7, 8^, these data show that CRB-2131 suppresses hippocampal AD-related neuroinflammation.

The above studies were all conducted with 3.5-month-old 5XFAD mice, which is the age where 5XFAD mice start developing amyloid plaques and neurodegeneration. At 6 months, these phonotypes are well established. Our experiments above showed that 10 weeks of prophylactic CRB-2131 treatment prevented these mice from developing brain ROS, lipid peroxidation, phosphorylated Tau, and cognitive problems that would normally be seen at 6 months. To test whether CRB-2131 can also treat established disease, we orally treated 6-month-old 5XFAD mice with 10 weeks of 10 mpk CRB-2131 (*q.d.*, and *P.O.*) (Fig. S4B) and then measured the levels of pTau, GFAP^+^ astrocytes and Iba-1^+^ microglia in the DG of the hippocampus. This therapeutic CRB-2131 treatment suppressed the levels of pTau in the cortex and the hippocampus, GFAP^+^ astrocytes and Iba-1^+^ microglia in the DG of the hippocampus, compared to vehicle-treated old 5XFAD mice (Fig. 3B and Fig. S6A to S6D). Interestingly, the therapeutic effect of CRB-2131 on the regulation of pTau, GFAP^+^ astrocytes and Iba-1^+^ microglia showed better than the positive control donepezil (6-month-old mice, 10 μg/kg, *q.d*., *P.O.*, and 10 weeks) (Fig. 3B and Sig. S6A to S6D).

### Administration of CRB-2131 ameliorates learning and memory of 5XFAD mice

To examine the effect of CRB2131 on 5XFAD mouse cognition, untreated WT, vehicle-treated 5XFAD, and prophylactically-treated 5XFAD mice (1, 3, or 10 mpk, *q.d*., *P.O*., and 10 weeks) underwent the novel object recognition (NOR), Y-maze, and Morris water maze (MWM) tests, which assess short-term memory, working memory, and spatial memory^28, 29, 30^. The movement of the mice was tracked with an automated video-tracking system (San Diego Instruments, San Diego, CA, USA). The NOR test examines short-term recognition memory^28^. The vehicle-treated 5XFAD mice did not display more curiosity with the novel object B in the second stage, which confirms the loss of cognition in these mice. By contrast, the 5XFAD mice that had been treated prophylactically with CRB-2131 showed similar curiosity towards B as the WT mice did (Fig. 4A). These results indicate that the Nox inhibitor compound ameliorate short-term recognition memory in 5XFAD mice.

**Figure 4.**
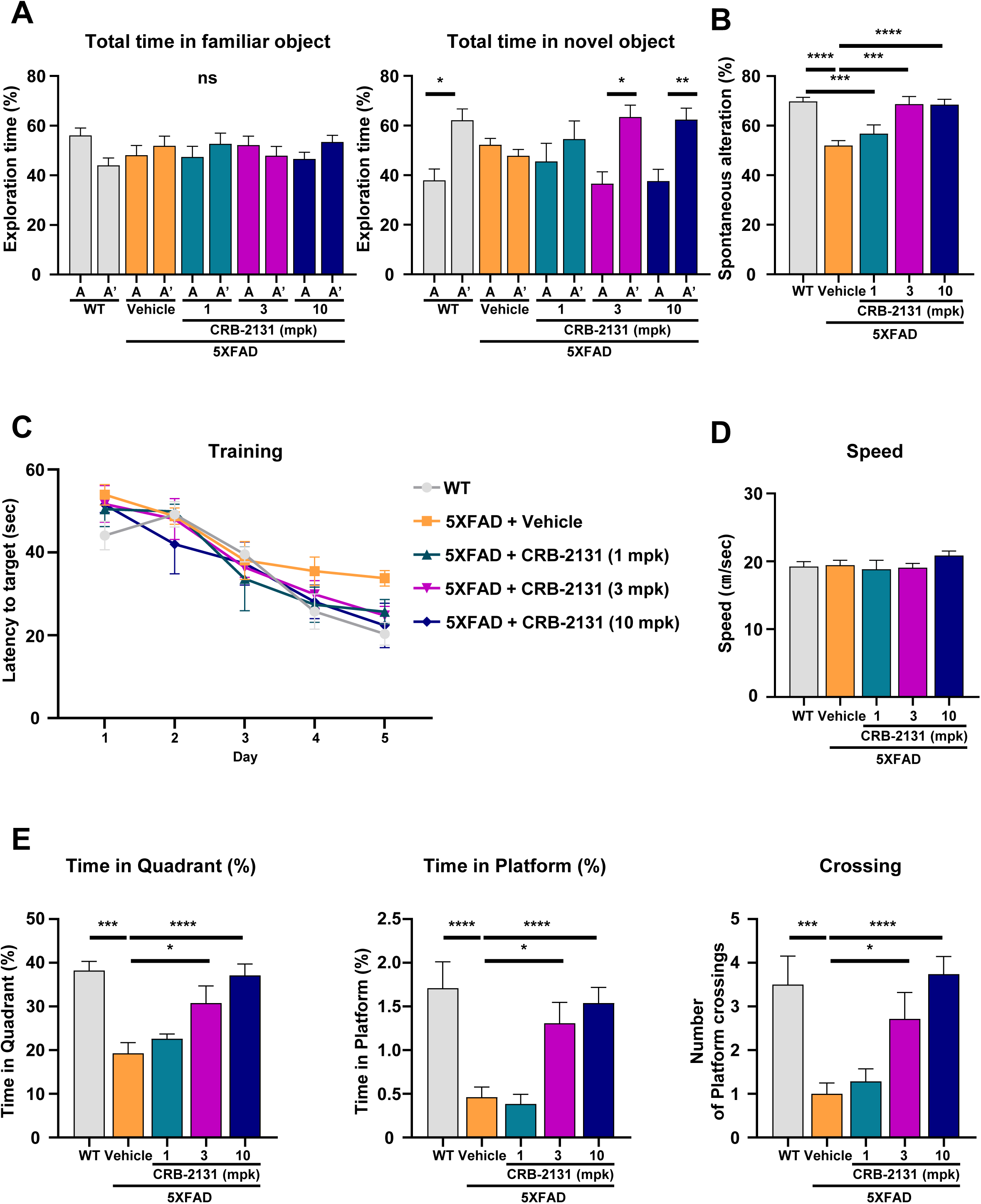
Prophylactic CRB-2131 improves behavioral-test outcomes in 5XFAD mice. Untreated WT, vehicle-treated 5XFAD, and prophylactic CRB-2131-treated 5XFAD mice (3.5-month-old mice, 1, 3, or 10 mg/kg of CRB-2131, *q.d.*, *P.O.*, and 10 weeks) were subjected to cognition tests. The mpk means mg/kg. (**A**) Novel object recognition (NOR) test. The time spent by the mice exploring the two same-shaped objects (A and A*) in the familiarization period and the familiar (A) and novel (B) objects in the second stage was quantified. (**B**) Y-maze test. The frequency of mouse explorations that involved sequentially investigating each of the three arms of the Y-shaped box (termed spontaneous alteration) was measured. (**C-E)** Morris water maze (MWM) test. **(C)** The mice underwent 5 consecutive days of training (4 trials per day) with a fixed platform. (**D-E)** The platform was removed on the sixth day, and the following were measured: speed with which the mice swam to the place the platform used to be (D), the time spent in the platform quadrant (E, left), the time spent at the place the platform used to be (E, middle), and the number of times the mice swam over the place the platform used to be (E, right). All quantitative data in this figure are shown as mean ± SD. *p<0.05, **p<0.01, ***p<0.005, ****p<0.0001, as determined by one-way ANOVA followed.

The Y-maze test examines short-term spatial memory^29^. In this test, the mice are placed in a Y-shaped box, which causes normal mice to investigate each arm in a sequential manner termed spontaneous alteration. In untreated WT mice, 70% of the exploration over 8 min involved spontaneous alterations. This was reduced to 50% in the vehicle-treated 5XFAD mice, but prophylactic CRB-2131 treatment caused full recovery of the spontaneous alteration in the 5XFAD mice (Fig. 4B). It indicates that the Nox inhibitor CRB-2131 effectively improves short-term spatial memory in 5XFAD mice.

The MWM test measures spatial learning and memory^30^. With regard to latency to target, the untreated WT mice showed steady improvement over the 5 training days. This was less marked in the vehicle-treated 5XFAD mice but the 5XFAD mice with prophylactic CRB-2131 treatment demonstrated similar latency as the WT mice (Fig. 4C). With regard to the variables measured after the platform was removed, the vehicle-treated 5XFAD mice demonstrated similar swimming speeds as the WT mice but they struggled to locate the target quadrant, as shown by less time spent in the southwestern quadrant and at the place the platform used to be and fewer crossing over the place the platform used to be. By contrast, the 3 mpk and especially 10 mpk CRB-2313-treated 5xFAD mice resembled the WT mice in all variables (Fig. 4, D and E). Thus, prophylactic CRB-2313 treatment improves spatial learning and memory in 5XFAD mice.

We then compared the ability of CRB-2131 to protect 5XFAD mice (3.5-month-old) from cognitive damage to donepezil, which is a conventional cholinesterase inhibitor therapy for AD^31^. CRB-2131 at 10 mpk protected the cognition of the 5XFAD mice better than 10 μg/kg donepezil given in the same regimen (*q.d.*, *P.O.*, and 10 weeks), as shown by the NOR, Y-maze, and MWM tests (Fig. S7).

To test whether CRB-2131 can also treat established disease, we orally treated 6-month-old 5XFAD mice with 10 weeks of 10 mpk CRB-2131 QD (Fig. S4B) and subjected the 8.5-month-old mice to the NOR, Y-maze, and MWM tests. This therapeutic CRB-2131 treatment improved the cognition of the 5XFAD mice, often almost to the level seen in the untreated WT mice (Fig. 5A to 5F). This treatment also performed better than the positive control donepezil (6-month-old mice, 10 μg/kg, *q.d.*, *P.O.*, and 10 weeks) (Fig. 5A to 5F).

**Figure 5.**
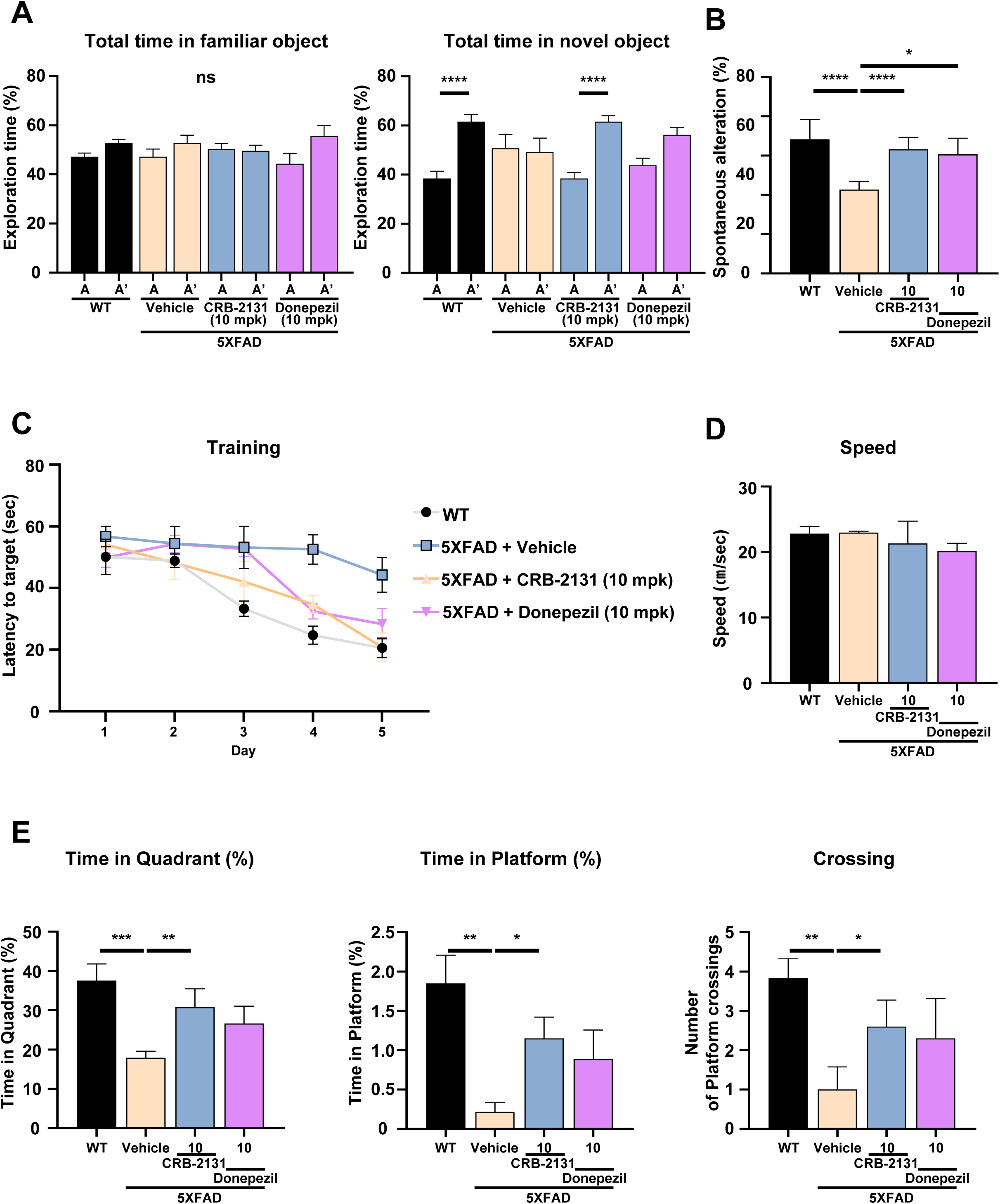
Therapeutic CRB-2131 improves behavioral-test outcomes in 5XFAD mice. Untreated WT, vehicle-treated 5XFAD, and prophylactic CRB-2131-treated 5XFAD mice (6.0-month-old mice, 10 mg/kg of CRB-2131, *q.d.*, *P.O.*, and 10 weeks) were subjected to cognition tests. The mpk means mg/kg. (**A**) Novel object recognition (NOR) test. The time spent by the mice exploring the two same-shaped objects (A and A*) in the familiarization period and the familiar (A) and novel (B) objects in the second stage was quantified. (**B**) Y-maze test. The frequency of mouse explorations that involved sequentially investigating each of the three arms of the Y-shaped box (termed spontaneous alteration) was measured. (**C-E)** Morris water maze (MWM) test. **(C)** The mice underwent 5 consecutive days of training (4 trials per day) with a fixed platform. (**D-E)** The platform was removed on the sixth day, and the following were measured: speed with which the mice swam to the place the platform used to be (D), the number of times the mice swam over the place the platform used to be (E, left), the time spent in the platform quadrant (E, middle), and the number of times the mice swam over the place the platform used to be (E, right). All quantitative data in this figure are shown as mean ± SD. *p<0.05, **p<0.01, ***p<0.005, ****p<0.0001, as determined by one-way ANOVA followed.

### CRB-2131 protects against neuronal death and promotes neurogenesis in 5XFAD mice

To further determine the in vivo protective mechanism of CRB-2131, we asked whether it could prevent neuronal-cell death: Aβ is well known to drive neuronal-cell death in AD^32, 33^. Thus, 5XFAD mice were treated prophylactic CRB-2131 treatment (3.5-month-old mice, 1, 3, or 10 mpk, *q.d.*, *P.O.*, and 10 weeks), after which the brain was subjected to immunohistochemistry with anti-NeuN^+^ as a marker of mature neuron. The loss of mature neurons (NeuN^+^) in the DG of hippocampus in 5XFAD mice model was attenuated by CRB-2131 in a dose-dependent manner (Fig. 6A and Fig. S8A). To investigate the protective effect of CRB-2131 on mature neuronal death in the DG of the hippocampus in 5XFAD mice (3.5-month-old), we measured co-localization between NeuN^+^ and TUNEL staining. Vehicle-treated 5XFAD mice increased cell death of mature neuron (TUNEL^+^NeuN^+^) in the DG of the hippocampus. However, this was reversed by prophylactic CRB-2131 treatment (Fig. 6B and 6C).

**Figure 6.**
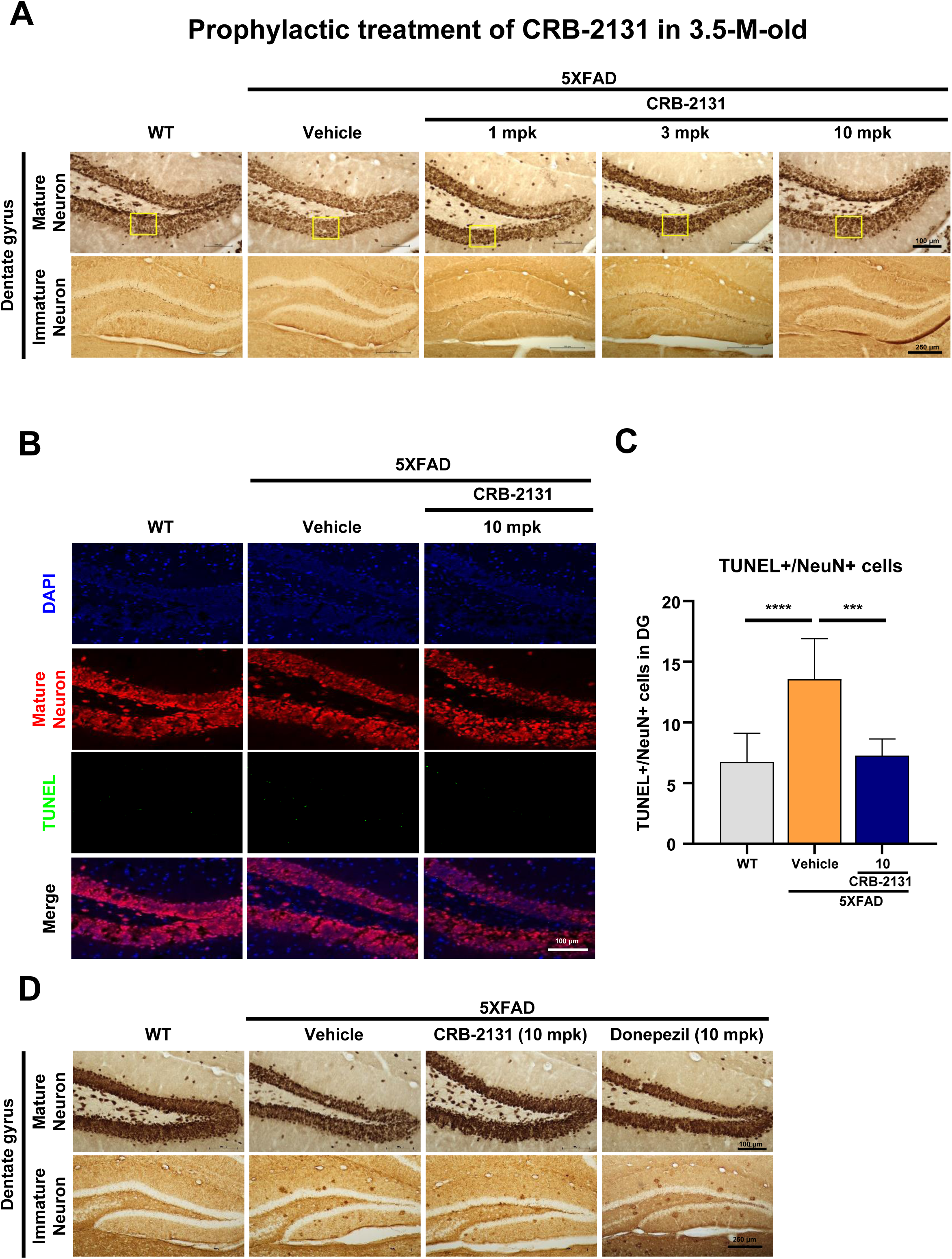
Prophylactic and therapeutic CRB-2131 protects 5XFAD mice from mature neuron death and enhances the regeneration of immature neurons. (**A**) Prophylactic CRB-2131-treated 5XFAD mice (3.5-month-old mice, 1, 3, or 10 mg/kg of CRB-2131, *q.d.*, *P.O.*, and 10 weeks) were subjected to immunohistochemistry to determine the mature neurons (NeuN^+^) and immature neuron (DCX^+^) in the DG. The TUNEL staining of the mature neurons was also assessed. Representative images of NeuN and DCX staining. The mpk means mg/kg. (**B**) Co-localization of NeuN expression and TUNEL staining in the hippocampus (10 mg/kg CRB-2131). (**C**) Quantification of the mature (NeuN^+^) TUNEL-stained hippocampal neurons. (**D**) Therapeutic CRB-2131-treated 5XFAD mice (6.0-month-old mice, qd, P.O., 10 weeks) were subjected to immunohistochemistry to determine the mature neurons (NeuN^+^) and immature neuron (DCX^+^) in the DG. The TUNEL staining of the mature neurons was also assessed. Representative images of NeuN and DCX staining. All quantitative data in this figure are shown as mean ± SD. *p<0.05, **p<0.01, ***p<0.005, ****p<0.0001, as determined by one-way ANOVA followed.

The hippocampus undergoes adult hippocampal neurogenesis, which produces new neurons that are then continuously incorporated into the DG^34, 35^. The hippocampus is one of the most affected brain regions in AD because of loss of adult hippocampal neurogenesis^34, 35^. This was observed in 5XFAD mice^34, 35^. To test whether CRB-2131 can shape adult hippocampal neurogenesis, 5XFAD mice were treated prophylactically with CRB-2131 (3.5-month-old mice, 1, 3, or 10 mpk, *q.d.*, *P.O.*, and 10 weeks) and then underwent brain immunohistochemistry to measure the doublecortin-expressing (DCX^+^) immature neurons in the DG. The vehicle-treated 5XFAD mice demonstrated decreased DCX^+^ neuron numbers, and this was reversed by CRB-2131 in a dose-dependent fashion (Fig. 6A and Fig. S8B). Thus, CRB-2131 enhanced the regeneration of immature neurons.

To validate the in vivo therapeutic activity of CRB-2131 in old 5XFAD mice (6-month-old mice, 10 mpk CRB-2131, *q.d.*, *P.O.*, and 10 weeks), we questioned whether it could prevent neuronal cell death and stimulate immature neuronal regeneration. Therapeutic treatment of CRB-2131 showed suppression of mature neuronal cell death and induction of immature neuronal regeneration (Fig. 6D and Fig. S8C and S8D).

To confirm these findings, untreated WT, vehicle-treated 5XFAD, and therapeutically-treated 5XFAD mice (3.5-month-old mice, 1, 3, or 10 mpk CRB-2131, *q.d.*, *P.O.*, and 10 weeks) were injected on four consecutive days with BrdU at 5 months of age (Fig. S4). BrdU is a marker of newly generated cells in the brain. Immunohistochemistry of the hippocampus showed co-localization of NeuN^+^ mature neurons and DCX^+^ immature neurons with BrdU in the WT mice, but this was suppressed in vehicle-treated 5XFAD mice. However, CRB-2131-treated 5XFAD mice showed restored co-localization of NeuN^+^ and DCX^+^ with BrdU (Fig. 7, A and B and fig. S9, A and B). Interestingly, this was observed when 6.0-month-old WT and vehicle- or CRB-2131-treated 5XFAD mice (therapeutic regimen) were injected with BrdU at 7.5 months of age. CRB-2131 restored BrdU co-localization with NeuN^+^ or DCX^+^ in old 5XFAD mice (Fig. 7, C and D and Fig. S10A and S10B). These findings suggest that by suppressing tauopathy and neuroinflammation, the Nox inhibitor CRB-2131 promotes an environment that reduces neuronal death and promotes immature-neuron regeneration. Thus, CRB-2131 induces a more resilient brain.

**Figure 7.**
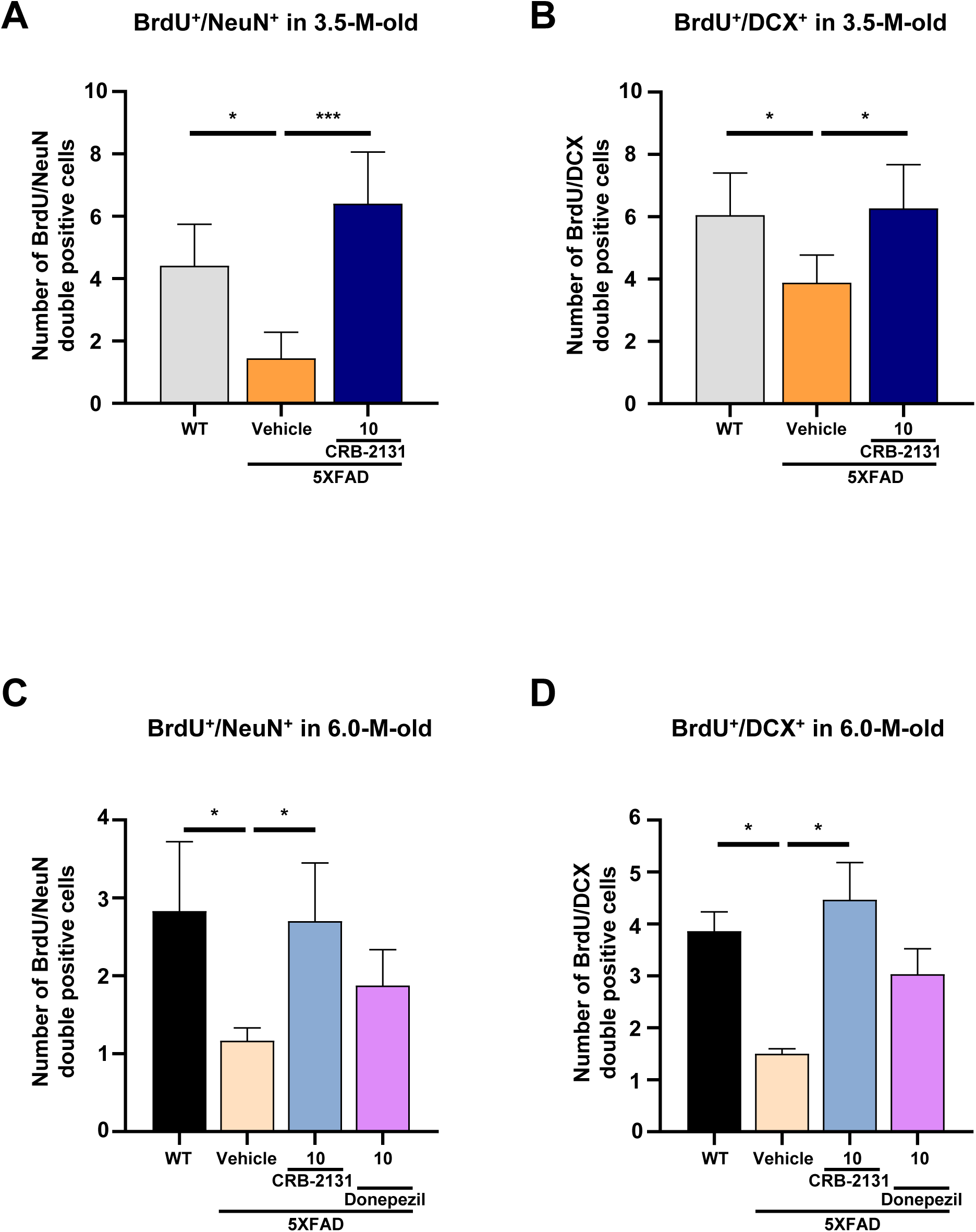
Prophylactic and therapeutic CRB-2131 promotes neurogenesis, as shown by colocalization of BrdU with mature and immature neurons. (A-B) Colocalization of BrdU with mature neuron (NeuN) and immature neuron (DCX^+^) in DG of 5XFAD mice (3.5-month-old mice, *q.d.*, *P.O.*, and 10 weeks) with or without CRB-2131 administration (10 mg/kg). The mpk means mg/kg. (**A)** Colocalization of BrdU with mature neuron (NeuN) in DG of 5XFAD mice with or without CRB-2131 administration (10 mg/kg). Quantification of mature neuron (NeuN) and BrdU staining of hippocampus in 5XFAD mice with or without CRB-2131 administration (10 mg/kg). **(B)** Colocalization of BrdU with immature neuron (DCX^+^) in DG of 5XFAD mice with or without CRB-2131 administration (10 mg/kg). Quantification of immature neuron (DCX^+^) and BrdU staining of hippocampus in 5XFAD mice with or without CRB-2131 administration (10 mg/kg). **(C-D)** Colocalization of BrdU with mature neuron (NeuN) and immature neuron (DCX^+^) in DG of 5XFAD mice (6.0-month-old mice, *q.d.*, *P.O.*, and 10 weeks) with or without CRB-2131 administration (10 mg/kg). (**C)** Colocalization of BrdU with mature neuron (NeuN) in DG of 5XFAD mice with or without CRB-2131 administration (10 mg/kg). Quantification of mature neuron (NeuN) and BrdU staining of hippocampus in 5XFAD mice with or without CRB-2131 administration (10 mg/kg). **(D)** Colocalization of BrdU with immature neuron (DCX^+^) in DG of 5XFAD mice with or without CRB-2131 administration (10 mg/kg). Quantification of immature neuron (DCX^+^) and BrdU staining of hippocampus in 5XFAD mice with or without CRB-2131 administration (10 mg/kg). All quantitative data in this figure are shown as mean ± SD. *p<0.05, ***p<0.005, as determined by one-way ANOVA followed.

### PET/CT confirms that CRB-2131 promotes neuronal regeneration in 5XFAD mice

Positron Emission Tomograph/Computed tomography (PET/CT) is a key molecular-imaging tool. Since AD patients typically exhibit decreased neuronal activity, particularly in the temporal and parietal cortex^36, 37^. In particular, ^18^F-fluorodeoxyglucose (FDG)-PET is used to evaluate neuronal glucose metabolism, since it primarily reflects neuronal connectivity. Since we found that CRB-2131 stimulates hippocampal neurogenesis in 5XFAD mice (Figs. 6 and 7), we subjected 3.5-month-old WT and vehicle- or CRB-2131-treated 5XFAD mice (prophylactic regimen, 10 mpk, *q.d.*, *P.O.*, and 10 weeks) to [^18^F]FDG-PET and CT brain imaging. The vehicle-treated 5XFAD mice had lower glucose-corrected standardized uptake values (SUVglc) in the whole brain than age-matched WT mice (Fig. 8A). This was reversed in the CRB-2131-treated 5XFAD mice (Fig. 8B). Moreover, when the mice were compared in terms of functional connectivity, the lower functional connectivity in the vehicle-treated 5XFAD mice was restored to normal levels in the prophylactic CRB-2131-treated 5XFAD mice. Notably, one of these nodes included the occipital and parietal temporal cortex (Fig 8C, and Fig. S11). Thus, PET/CT supports the notion that CRB-2131 stimulates neuronal regeneration, enhancing neuronal activity. This suggests that the Nox inhibitor CRB-2131 could be a novel therapeutic candidate for AD.

**Figure 8.**
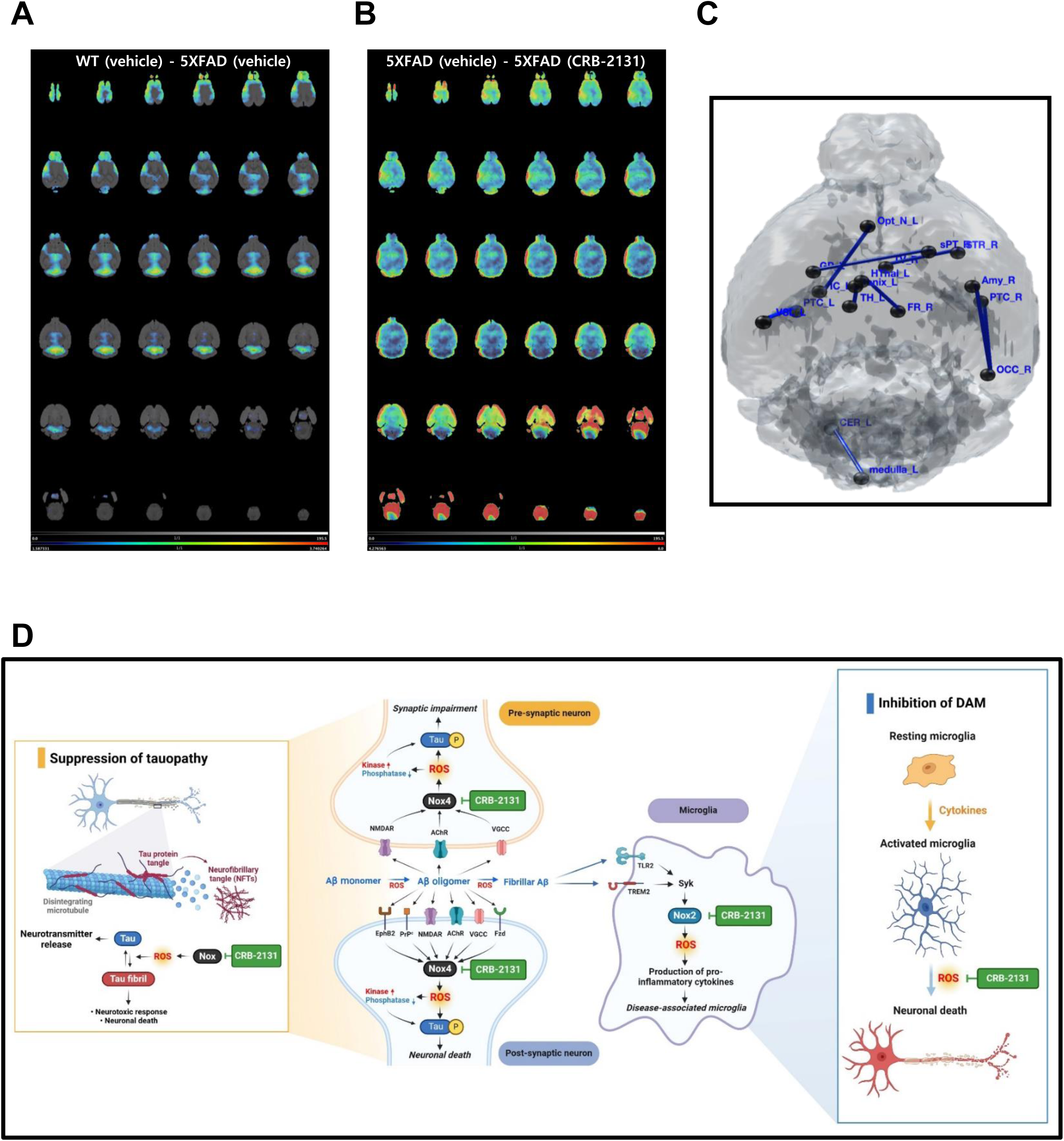
PET/CT testing for neuronal regeneration in prophylactic CRB-2131-treated 5XFAD. Prophylactic CRB-2131-treated 5XFAD mice (3.5-month-old mice, *q.d.*, *P.O.*, and 10 weeks) underwent PET/CT at 6 months of age. (**A** and **B**) Ratio of SUVglc in (A) control vehicle-treated 5XFAD mice relative to WT mice and (B) vehicle-treated 5XFAD mice relative to CRB-2131-treated 5XFAD mice. **C)** Functional connectivity between vehicle-treated and CRB-2131-treated 5XFAD mice. (**D**) Proposed model by which CRB-2131 improves cognitive ability in the 5XFAD model of AD. Specifically, by suppressing Nox-induced oxidative stress, CRB-2131 reduces Tau hyperphosphorylation and inhibits neuroinflammatory pathways, thereby facilitating the regeneration and repair of the neural tissues.

## Discussion

Since reactive oxygen species (ROS) are known to be toxic, a balance between ROS generation and elimination in cells is tightly regulated. In a pathological stage such as AD, activation of various ROS generation system stimulates uncontrolled ROS generation^38, 39, 40^. With the balance of ROS homeostasis is disrupted, the levels of ROS gradually increase in the brain, closely linked to the pathogenesis of AD. It is well known that the brain is highly oxygen- and energy-demanding organ, making it particularly susceptible to oxidative stress. This vulnerability is due to its elevated levels of metabolic enzymes that contain redox-related metal ions such as iron and copper and abundance of lipids that are highly prone to ROS-induced oxidization. Moreover, while the blood-brain barrier shields the brain tissue from external influences, it also intensifies the impact of events occurring within it, such as ROS accumulation. The resulting high ROS levels in the brain are closely associated with AD.

Excessive ROS production caused by Aβ-induced Nox activation in brain tissue can disrupt redox balance and lead to toxic conditions such as tauopathy and neuroinflammation. Recent findings underscore the pivotal role of tau pathology in the pathogenesis of dementia and suggest that modulating excessive reactive oxygen species (ROS) production, driven by Aβ-induced NADPH oxidase (Nox) activation, may be crucial in mitigating the progression of tauopathy ^10^. Therefore, effectively regulating ROS production through Nox inhibitor can positively influence physiological processes in the brain tissue. In the brain tissue of AD patients, reduced neurogenesis in the hippocampus leads to a decrease in immature neurons. Essentially, in aging and in the brain tissue of AD patients, elevated levels of ROS create an unfavorable environment, resulting in a decline in neurogenesis. Under these adverse conditions, decreasing ROS through Nox inhibitor CRB-2131 treatment can potentially shift the detrimental environment, marked by tauopathy and neuroinflammation, into a more supportive one for neurogenesis to thrive.

The close relationship between ROS and AD has led to the development and clinical trialing of numerous dietary antioxidants (e.g. vitamin C) as AD therapies^41, 42^. However, the trial results have been highly inconsistent and many trials have failed to show improvements in AD patients. The main reason for this may be that dietary antioxidant supplementation does not sufficiently eliminate the ROS in the brain of AD patients because the antioxidants become distributed throughout the entire body rather than concentrating in the brain. Thus, a better approach would be to target the enzymes that regulate antioxidant levels in the brain. Hitherto, this more fundamental approach to AD pathology has not been as strongly researched as dietary antioxidants.

A possible target of the latter approach could be Nox: its only function is to produce ROS, it is responsible for considerable ROS generation, and the progression of disorders such as AD is strongly linked to the overexpression of Nox isozymes in the brain. Thus, Nox inhibitors could specifically reduce the high ROS levels in the brain of patients with AD. We show here that a new candidate Nox inhibitor, CRB-2131, effectively protects 5XFAD AD-model mice from cognitive impairment, both when it is administered before the disease manifests itself and after the disease is well established. Specifically, CRB-2131 markedly reduces the ROS, lipid peroxidation, and phospho-Tau levels in the DG and improves the cognition of the mice when performing various behavioral tests. This associates with decreased neuroinflammation, as shown by fewer GFAP^+^ astrocytes and Iba-1^+^ microglia in the DG. Significantly, it also associates with less mature neuron death and more regeneration of immature neurons via adult hippocampal neurogenesis (Figs. 6 and 7).

Fibrillary Aβ stimulates various receptors and channels including NMDAR, AChR, EphB2, and VGCC in neurons^43, 44, 45, 46, 47^. The binding of these receptors with Aβ aggregates can induce ROS generation through Nox1/4, predominant isozymes in neurons. ROS can inhibit protein tyrosine phosphatase (PTPase) by oxidizing the thiolate anion in its active center to sulfenic acid (-SOH). This oxidation events can disrupt the balance between PTK and PTPase^48, 49^, thus relatively enhancing PTK activity and increasing tyrosine phosphorylation of cell signaling-related proteins such as Tau. The phosphorylation of Tau prompts Tau to dissociate from microtubule, causing microtubule fibers to break down. This in turn associates strongly with synaptic impairment and neuronal death. We observed that Aβ- and Nox1/4 cascade-dependent ROS generation stimulate Tau phosphorylation in neuronal cells, and that Tau aggregation in neurofibrillary tangles was increased in 5XFAD mice^4, 5, 6^. The Nox inhibitor CRB-2131 compound suppressed Aβ-induced Tau phosphorylation in neuronal cells. It also reduced Tau phosphorylation and neurofibrillary tangles in 5XFAD mice. Thus, the Nox inhibitor CRB-2131 alleviates tauopathy in a mouse model of AD (Fig. 8D).

The non-receptor tyrosine kinase called spleen tyrosine kinase (SYK) is expressed in various cell types and is coupled with immune receptors such as the B-cell receptor (BCR), T-cell receptor (TCR), Toll-like receptor 4 (TLR4), and triggering receptor expressed on myeloid cells-2 (TREM2)^50, 51, 52^. TREM2 is a well-known receptor of fibrillary Aβ and associates closely with AD. TREM2 on microglia can recognize fibrillary Aβ, which then activates SYK. Previous reports show that SYK-deficient microglia fail to accumulate Aβ plaques, which decreases brain pathology and behavioral deficits. Astrocytes and microglial cells exclusively express Nox2 isozyme. We showed in the present study that Aβ stimulation enhanced microglial-cell migration and their production of pro-inflammatory cytokines, including TNF-α and IL-1β, and that these neuroinflammatory events were suppressed by treatment with CRB-2131. The fact that Aβ-dependent neuroinflammation can be regulated by CRB-2131 highlights the potential of Nox inhibitors as novel therapeutics for treating AD (Fig. 3A).

We found that CRB-2131 inhibited all Nox isoforms. Several lines of evidence show that multiple Nox isozymes may participate in A. For example, high Nox1, Nox2, Nox3, and Nox4 levels are observed in the brain of AD patients and animal model of AD. Moreoever, transgenic mice lacking the catalytic subunit of Nox2 failed to develop oxidative damage, neurovascular dysfunction, and cognitive deficits despite the lack of Aβ plaques. In addition, neuronal knockdown of Nox4 ameliorated tauopathy and cognitive decline in a humanized mouse model of tauopathy. Furthermore, inhibiting or deleting Nox2 decreases inflammation-elicited neuronal production of ROS in vitro^17^ while deleting or inhibiting Nox2 reduces Aβ-induced microglial production of ROS and IL-1β in vitro ^19^. These observations suggest that a pan-Nox inhibitor such as CBR-2131 may have better potential as a treatment for AD than more specific Nox inhibitors.

AD associates with reduced neurogenesis in the hippocampus, which decreases immature neurons, thereby impairing neuronal regeneration. We found that CRB-2131 not only prevented this in 5XFAD mice developing AD, it also reversed it in 5XFAD mice that have established AD. It also reduced mature neuron apoptosis in a prophylactic and therapeutic fashion. Thus, by suppressing brain ROS and the resulting tauopathy and neuroinflammation, the Nox inhibitor CRB-2131 treatment can create a brain environment that supports neurogenesis and reduces neuronal death.

PET/CT analysis of neural activity and connectivity, particularly in the temporal and parietal cortex, is used to diagnose AD^36, 37, 53^. Our PET/CT analysis of vehicle- and prophylactic CRB-2131-treated 5XFAD mice showed that CRB-2131 improved both neural activity and connectivity. Notably, connectivity between the occipital and parietal temporal cortex was also improved by CRB-2131. This is significant because interactions between the occipital cortex and other areas such as parietal cortex support visuospatial working memory. Moreover, it has been shown that improved occipital cortex function correlates with enhanced visuospatial memory and stimulating the occipital cortex with transcranial direct current stimulation boosts visual working memory, thus potentially enhancing visuospatial memory function ^54, 55^. Notably, we found that CRB-2131 strongly improved the performance of 5XFAD mice in the MWM test, which measures visuospatial memory. Thus, this improvement may correlate with the greater neural connectivity that was observed with PET/CT. Further studies on the role of neural connectivity with the occipital cortex in cognitive functions such as visuospatial memory are needed.

In summary, our research shows that the novel Nox inhibitor CRB-2131 effectively downregulates brain ROS levels, Tau phosphorylation, and neuroinflammation in 5XFAD mice. This prevents mature neuronal-cell death and promotes neuronal regeneration, ultimately generating brain resilience that protects the mice from learning and memory deficits. These effects were observed both before and after brain pathology emerged. Thus, the novel Nox inhibitor CRB-2131 shows potential as a new therapeutic agent for AD.

## Methods

### Reagents

Anti-Phospho-Tau (Ser202, Thr205) antibody was obtained from Invitrogen and anti-Iba1 antibody and anti-GFAP antibody were purchased from Wako. Anti-DCX antibody and anti-NeuN antibody were obtained from Thermo. Anti-BrdU antibody was purchased from Abcam and anti-4-hydroxynonenal (4-HNE) antibody was obtained from JaICA (Shizuoka, Japan).

### Animals

All animal procedures were approved by the Institutional Animal Care and Use Committee (IACUC) at Ewha Womans University. Mice were housed in specific pathogen-free environment under a 12 hrs light and/or dark cycle. Animal protocols including food and water were in compliance with NIH Guideline for the Care and Use of Laboratory Animals and have been approved by the Institutional Animal Care and Use Committee (IACUC) of Center for Laboratory Animal Sciences, Ewha Industry-University Cooperation Foundation, Ewha Womans University. All efforts were made to keep animal usage to a minimum, and male mice were used. 5XFAD (Tg 6799) breeding pairs were acquired from the Mutant Mouse Resource and Research Center (MMRRC) (Jax 034848) and crossed with C57BL/6 J mice to generate offspring for this study.

### Chemical synthesis of 4-Fluoro-N-(5-phenyl-1,3,4-oxadiazol-2-yl)-3-(trifluoromethyl) benzamide (CRB-2131)

All reagents were purchased from commercial vendors and used without further purification. The reaction was monitored by thin-layer chromatography (TLC) under UV light (254 nm). The solution of 5-phenyl-1,3,4-oxadiazole-2-amine (2.0 g, 12.41 mmol) in pyridine (50 mL) was added dropwise 4-fluoro-3-(trifluoromethyl)benzoyl chloride (2.25 mL, 14.89 mmol) and the reaction mixture was stirred at 73℃ overnight. After completion of the reaction, the mixture was cooled to room temperature, and then diluted with a 10% aqueous solution of HCl. The aqueous layer was extracted with EtOAc. The organic phase was washed with a 10% aqueous solution of HCl and water, dried over anhydrous MgSO4, and evaported at 50 ℃ to remove residual pyridine and solvents. The crude product was purified by solidification and trituration (CH2Cl2/MeOH/hexanes) to give the title compound CRB-2131 (85.3%, 3.72 g) as a white solid (mp 242.7 ℃); ^1^H-NMR spectra were obtained on a VNMRS 500 (Varian, USA), and 19F-NMR spectra were obtained on an AVANCE III HD 500 (Bruker, Germany) spectrometer at the KBSI Metropolitan Seoul Center, using DMSO-d₆ as the solvent. Chemical shifts were given in ppm using TMS as the internal standard, coupling constants are given in Hz. High resolution mass spectra (HRMS) were recorded on a Waters Xevo G2 Q-TOF mass spectrometer (Waters, MA, USA). ^1^H-NMR (500 MHz, DMSO-d6) δ 12.74 (s, 1H), 8.46 (d, J = 6.1 Hz, 1H), 8.41 (s, 1H), 7.97 (d, J = 6.6 Hz, 2H), 7.73 (t, J = 9.5 Hz, 1H), 7.63 – 7.61 (m, 3H); 19F-NMR (470 MHz, DMSO-d6) δ -59.07 (d, *J*= 12.2 Hz), -108.45 (m); HRMS (ESI) m/z: [M+H] + calcd for C16H10F4N3O2: 352.0709; found, 352.0699; purity ≥99% (as determined by RP-HPLC, method A, tR = 3.15 min; method B, tR = 4.39 min). The melting point was checked using melting point apparatus with USB port and MeltView software system MPA 100 (Stand ford Research Systems). The chemical purity of CRB-2131 submitted for biological testing was detected using a Waters HPLC system equipped with an autosampler and a PDA detector. HPLC method. Column: Fortis C18 column (150 mm × 4.6 mm, 5 μm packing diameter); flow rate: 1 mL/min; run time: 10 min; UV absorption: 254 nm; mobile phase A: 0.1% formic acid in water, and mobile phase B: 0.1% formic acid in CH3CN. Method A: Isocratic elution; 80% B; Method B: gradient elution; 70% to 80% B.

### Novel object recognition test

White acryl box (42 cm × 42 cm × 42 cm) and two kinds of objects with different shapes were used for the novel object recognition task. During the behavior test, room brightness was dimmed to reduce anxiety. All behaviors were recorded via a video camera and then analyzed (Harvard Apparatus, Holliston, MA, USA). During the familiarization phase, all mice were freely exposed to two of the same objects for 10 min. The 2 hours after familiarization, one object was kept as a familiar object, while another object was replaced with a novel object. All mice were also freely exposed to two of the different objects for 10 min. The mice were placed back in their home cage during the interval. The percentage of time spent interacting with A and A* in the familarization stage and with A and B in the second stage were recorded. In this test, each mouse is allowed to freely investigate two identical objects (denoted A and A*) placed in opposite corners of a box. After 10 min, it is returned to its cage for 3 h. A* is then replaced with a differently shaped object (denoted B) and the mouse is returned to the box for another 10 min. The time spent exploring A and A* in the first stage and A and B in the second stage is measured. The untreated WT mice spent equivalent time exploring A and A* in the first stage but spent less time exploring A and more time exploring B in the second stage. Thus, the experimental setup was effective. At the end of each trial, the bedding on the floor was changed and the apparatus was cleaned with 70% ethanol.

### Y-maze

To investigate cognition and spatial memory, spontaneous alternation test in the Y-maze was investigated in the mice. We performed the Y-maze test 10 weeks after CRB-2131 treatment. The alternation performance was performed using an asymmetric Y-maze, consisting of three identical arms (15 cm high × 9 wide cm × 40 cm long), and constructed using white acrylic material. All animals were placed in the same arm of the Y-maze and allowed to explore freely for 8 min. Unimpaired animals will enter the three arms in an alternating pattern termed spontaneous alternation. All movement were recorded and calculated using a video camera. The frequency of spontaneous alternations was measured by counting the number of times all three arms were entered sequentially (Harvard Apparatus, Holliston, MA, USA).

### Morris water maze

We performed a water maze test 10 weeks after CRB-2131 treatment to verify the improvement in cognitive function by CRB-2131. A circular water pool (90 cm in diameter) was filled with water (23 ± 2 °C temperature), and the water was mixed with edible dye (Bright White Liqua-Gel; Chefmaster, CA, USA) to make it opaque so that the platform (10 cm in diameter) was not visible. Visual cues of different shapes were marked on the walls of the four quadrants, North (N), West (W), South (S), and East (E), of the pool. Water was filled up to 1 cm above the platform. The platform was placed at the center of the southeastern (SE) quadrant. Learning training to remember the platform was performed for 5 consecutive days, and probe tests were performed on the 6th day. All mice were trained to remember the platform for 5 days, four times a day at each of the four points on the circle, and the probe test was performed on the 6th day. The subjects were placed into four quadrants facing the wall and allowed 1 min to reach the platform. Mice that found the platform within 1 min were left on the platform for 10 s for learning, and those that did not be find it were placed on the platform for 10 s. The probe test was performed after removing the platform. The mice started on the wall of the quadrant zone opposite to the platform, and their movements for 1 min were recorded using a tracking system (Harvard Apparatus, Holliston, MA, USA). The four repeats on the training days were averaged. Four variables were calculated from the recordings: the Mean locomotion speed on day 6; the time spent in the southeastern quadrant on day 6; the time spent on day 6 at the place where the platform used to be; and the number of times on day 6 the mice crossed over the place where the platform used to be.

### Immunohistochemistry staining

Ten weeks after CRB-2131 treatments, the mice were anesthetized with intraperitoneal injection of 2.5% Avertin (2,2,2-tribromoethanol) and immediately perfused through the heart with 0.9% saline followed by 4% paraformaldehyde in PBS. Brains were excised, post-fixed in 4% paraformaldehyde at 4 °C for 3days, and incubated in 30% sucrose at 4°C until reaching equilibrium. The brains were embedded in O.C.T. compound blocks at − 80°C. Sequential 30-μm coronal sections were obtained with a cryostat (CM30 50S; Leica). Every seven section (210 μm apart) of the brain (Bregma − 0.94 to − 2.70 mm) was used for immunohistochemistry.

All sections were washed in PBS and incubated in blocking solution (PBS, 5% normal goat serum, 0.2% Triton X100) for 1 h at room temperature. The sections were incubated with primary antibodies in blocking solution overnight at 4°C. The following primary antibodies were used for immunohistochemistry Anti-Phospho-Tau (Ser202, Thr205) antibody (1:2000– 4000, Invitrogen) was used to detect phophorylated tau proteins, anti-Iba1 antibody (1: 1000, Wako) was used to detect microglia, anti-GFAP antibody (1: 1000, Wako) was used to detect astrocyte, anti-DCX antibody (1: 1000, Thermo) was used to detect immature neuron, anti-NeuN antibody (1: 1000, Thermo) was used to detect mature neuron, and anti-BrdU antibody (1: 1000, Abcam) was used to detect cell proliferation. Biotinylated goat anti-rabbit antibody (VECTOR) was used as a secondary antibody followed by ABC solution (VECTOR). Then the areas that are immuinoreactive turned brawn by incubation with 3-3′diaminobenzidine (DAB, Dako). Eight coronal sections of hippocampus from Bregma −0.94 mm to −2.70 mm were collected in every 210 μm to stain and analize using ImagePro pluse 7.0 software.

### BrdU assay

BrdU (Sigma, St. Louis, MO) was injected with intraperitoneal route at 100 mg/kg once daily for four days 4 weeks before sacrifice and detected with an anti-BrdU antibody (Abcam).

### DHE (Dihydroethidium) Assay

The frozen brain sections were stained with 10 μM DHE (D1168, Invitrogen) for 30 min at 37°C. To evaluate superoxide production with DHE, red fluorescence (585 nm) was measured by the LSM 880 Airyscan (Carl Zeiss Vision System)^56^.

### TUNEL assay

Apoptotic DNA fragmentation was detected using the TUNEL method 57. After incubation, the cells were subjected to TUNEL staining in accordance with the manufacturer’s instructions (In Situ Cell Death Detection Kit, Fluorescein, Roche 11684795910). Apoptotic cells were detected as localized bright green cells (positive cells) in a blue background using an Zeiss confocal 880 for confocal microscopy.

### 18F-fluorodeoxyglucose-Positron Emission Tomograph/Computed tomography (FDG PET/CT)

The FDG brain images for each mouse were spatially normalized to a standardized FDG PET template of the murine brain using PMOD software version 3.7 (PMOD Technologies). Post-registration to the template, a three-dimensional brain mask was applied to the images, effectively setting all voxel values outside the delineated brain region to zero. ach parametric image was spatially normalized to an MRI template of the mouse brain. Image analyses were conducted using SnPM (Wellcome Trust Centre for Neuroimaging). The registered images underwent statistical analysis employing a t-test. Comparisons between the test and vehicle groups were performed using nonparametric permutation tests. An uncorrected p-value threshold of < 0.005 was applied for statistical significance 53, 58.

### Functional correlation and brain network construction

To construct brain networks, we utilized 53 nodes, each corresponding to one of the 53 volumes of interest (VOIs). We extracted intensity-normalized FDG uptake values from the VOIs of each mouse. Utilizing these uptake values, we computed correlation coefficients. Pearson’s correlation coefficients (r) between each pair of VOIs were determined in an intersubject manner, resulting in a 53 × 53 correlation matrix for each experimental group. To assess statistical differences in interregional correlations between the groups, we employed a permutation test on all possible connections between nodes. The interregional correlation matrices for all groups were transformed to Z-scores via Fisher transformation. Labels were randomly reassigned and permuted 5000 times for each of the 53 VOIs, followed by the calculation of interregional correlation matrices and subsequent Fisher transformation. Type I error rates were derived by comparing the observed Z-score for each connection with the Z-scores obtained from the permuted data. Statistically significant differences in connections between the vehicle and test groups were determined using a significance threshold of p < 0.005 53, 58.

### The prophylactic and therapeutic treatment of CRB-2131 in 5XFAD mice

5XFAD mice were treated with CRB-2131 either prophylactically before the onset of AD-like symptoms at 3.5 weeks of age, or therapeutically after the onset of symptoms at 6 months of age^59, 60, 61^. In both cases, CRB-2131 was administered orally at 1, 3, or 10 mg/kg on a daily basis (q.d.) for 10 weeks. The prophylactically- and therapeutically-treated mice were thus sacrificed at 6 and 8.5 months of age, respectively. The mice underwent cognition tests or PET/CT scans 1 week before sacrifice. After sacrifice, immunohistochemistry and other experiments were conducted on the brain tissue.

## Statistical analysis

All data were expressed as mean ± standard error of the mean (SEM). All statistical analyses were performed using the GraphPad Prism 8 software (GraphPad Software Inc., San Diego, CA, USA). All results were statistically analyzed using one-way analysis of variance (ANOVA) followed by Tukey’s post-hoc test. As an exception, comparisons between groups over days in the MWM training were analyzed by two-way repeated measure ANOVA with Tukey post-hoc test was conducted. Data for Aβ quantification were analyzed using unpaired t-test (two-tailed). For all analyses, statistical significance was set at p < 0.05.

## Acknowledgements

We thank Professor E.S. Kim (School of Medicine, Yonsei University) for critical reading the manuscript. This research was supported by NRF grants (RS-2024-0039829) and (NRF-2013M3A9D5072560) from Korea Mouse Phenotyping Projects of the National Research Foundation (NRF) funded by the Ministry of Science and ICT, Republic of Korea and the Starting Growth Technological R&D Program (20165925) funded by the Ministry of SMEs and Startups (MSS, Korea).

## Author contributions

J.L. conceived and designed the experiments, interpreted the data, and wrote the manuscript. S.Y., H.E.L., D.U.J., J.M.S, and I.H.L. designed the experiments, interpreted the data. S.S, Y.J., and H.P. contributed to synthesis and provide Nox inhibitor and its derivatives. E.Y.B., S.J.K., and H-Y.L contributed to PET/CT data analysis. Y.C and Y.S.B supervised all experiments, interpreted data, and wrote the manuscript.

## Additional information

Additional supporting information can be found online in the Supporting Information section at the end of this article.

